# Multi-state occupancy model estimates probability of detection of an aquatic parasite using environmental DNA: *Pseudoloma neurophilia* in zebrafish aquaria

**DOI:** 10.1101/2022.02.16.480730

**Authors:** Corbin J. Schuster, Michael L. Kent, James Peterson, Justin L. Sanders

## Abstract

Detecting the presence of important parasites within a host and its environment is critical to understanding the dynamics that influence a pathogens ability to persist, while accurate detection is also essential for implementation of effective control strategies. *Pseudoloma neurophilia* is the most common pathogen reported in zebrafish (*Danio rerio*) research facilities. The only assays currently available for *P. neurophilia*, are through lethal sampling, often requiring euthanasia of the entire population for accurate estimates of prevalence in small populations. We present a non-lethal screening method to detect *Pseudoloma neurophilia* in tank water based on detection of environmental DNA (eDNA) from this microsporidum, using a previously developed qPCR assay that was adapted to the digital PCR (dPCR) platform. Using the generated dPCR data, a multi-state occupancy model was also implemented to predict the probability of detection in tank water under different flow regimes and pathogen prevalence. The occupancy model revealed that samples collected in static conditions were more informative than samples collected from flow-through conditions, with a probability of detection at 80% and 47%, respectively. There was also a positive correlation with the prevalence of infection in water and prevalence in fish based on qPCR.

## 1. Introduction

The zebrafish as a biomedical model is widely used in many studies ranging from immunological and infectious disease to developmental biology and neuro-behavioral studies. The ease of breeding and housing and the development of genetic tools has facilitated the expansion of this model into nearly every field of biology. Some genetic lines produced for specific experiments breed poorly and require labor intensive husbandry conditions to maintain even a small population, creating several challenges to the production of embryos needed to fulfill experimental protocols (Avdesh et al. 2012). A common threat to laboratory zebrafish and overall zebrafish facility operations is the presence of infectious diseases. While many aspects of the husbandry conditions of this animal are known, many remain unknown- or poorly understood, which may contribute to the negative impacts of pathogens. The presence of infectious diseases in laboratory zebrafish has significant impacts to the maintenance of zebrafish populations and may be a confounding factor in research results, potentially biasing the conclusions of many studies (Kent et al. 2012).

A microsporidian parasite, *Pseudoloma neurophilia*, is an ongoing threat to the zebrafish model. This parasite continues to be prevalent in zebrafish research facilities that report to the diagnostic service of the Zebrafish International Resource Center (ZIRC) in Eugene, Oregon (Murray et al. 2011, Kent et al. 2020a). *P. neurophilia* is an obligate intracellular parasite that causes chronic infections in zebrafish and infects a broad range of fishes (Sanders et al. 2016). Infections by *P. neurophilia* are largely asymptomatic. However, a subset of infected populations may present general clinical signs such as emaciation and skeletal deformities. Histologically, spores occur in the central nervous system and may cause associated gliosis (Spagnoli et al. 2015), whereas the parasite infects other organs (Sanders et al. 2014) causing other various forms of inflammation (myositis, menixitis, and encephalitis) that are also associated with the infection. Important to the integrity of the zebrafish model, *P. neurophilia* causes significant alterations in behavior and has been reported to alter transcripts in the brain, downregulating several genes involved in immune function. Moreover, infections cause reduced fecundity and growth, while stress exacerbates *P. neurophilia* prevalence in large populations (Ramsay et al. 2009).

The biology of the parasite promotes its ability to survive in the environment, as it develops into a hearty resistant spore that is highly resistant to disruption and even regular disinfection protocols (Ferguson et al. 2007). The development of *P. neurophilia* has three major phases: an infectious phase, which is free spores found in the environment, and two intracellular phases, proliferative and sporogonic. During the intracellular growth phases, several developmental stages occur, but it is free spores in the environment that can be a useful target for non-lethal diagnostics. Additionally, transmission occurs both vertically and horizontally, which creates numerous opportunities for detection of the parasite, as spores are released in the environment during spawning, from decomposing carcasses, and through urine and feces (Sanders et al. 2013).

Environmental DNA (eDNA) assays are commonly used by ecologists to detect and quantify organisms in water, air, and soil (Rees et al. 2014; Barnes et al. 2014; Bass et al. 2015). In terrestrial systems, parasites are often detected in the soil or feces (Almazan et al. 2001; Mandarino-Pereira et al. 2010; Nagamori et al. 2018;). Whereas analysis of feces is usually not practical for aquatic parasites, the water itself provides a useful medium for detecting parasites simply through filtration. Hence, surveillance in aquatic systems has been deployed for detection of several human and wildlife pathogens, including parasites (Berger & Aubin-Horth 2018; Sieber et al. 2020; Amarasiri et al. 2021). Notably, such have been developed for detecting common water-born human parasites, such as *Cryptosporidium* spp. and *Giardia lamblia* using filtration of water (Guy et al. 2003).

Water tests have also been developed for a variety of fish parasite taxa – e.g., myxozoans, helminths and protozoa. These include *Ceratonova shasta* (Hallett et al. 2012), *Nanophyetus salmonicola* (Purcell et al. 2017) *Gyrodactylus salaris* (Rusch et al. 2018), and *Dactylogyrus* spp. (Trujillo-González, 2019), *Neobenedenia girellae* (Agawa et al. 2016), and *Chilodonella hexasticha* (Bastos-Gomes et al. 2017). *Ichthyophthirius multifiliis* (Howell et al. 2019). Furthermore, Shea et al. (2020) used a multi-plex PCR assay to screen for a panel of salmon parasites in seawater. In the case of zebrafish parasites, an eDNA assay has recently been developed for the capillarid nematode *Pseudocapillaria tomentosa*. The assay for *P. tomentosa* was shown to effectively detect the pathogens in feces, detritus, and water samples (Norris et al. 2020). In contrast, *P. neurophilia* is not easily detectable in water with current assays (Sanders et al. 2013; Crim et al. 2017; Miller et al. 2019).

Detection by PCR is commonly used in eDNA methods, but there are some general limitations, particularly when the target species is present in low abundance, which may be compounded by the fact PCR is unable to distinguish between life stages of the parasite (Kralik et al. 2017). Often, quantitative PCR (qPCR) is utilized in the case of environmental testing, but due to advancements in technologies, such as the implementation of digital PCR (dPCR), the sensitivity of diagnostic assays has dramatically improved in recent years (Koepfli et al. 2016). Regarding pathogen detection in environmental samples, dPCR can be more sensitive than qPCR (Yang et al. 2014; Wilson et al. 2015; Norris et al. 2020) due to its inherent resistance to inhibition, and ability to provide more precise quantification without the need for a standard curve (Quan et al. 2018). Quantification of target DNA by dPCR is achieved by partitioning each sample into thousands of reactions, in such a fashion that each reaction is analyzed for the amplification product in an endpoint PCR and measured.

While molecular methods provide an avenue for detection, occupancy models enable the estimation of the proportion of an area occupied by an organism, when the target species is not detected with certainty (MacKenzie et al. 2004). Thus, these models are used to account for imperfect detection of organisms in surveys and to determine the probability of the true presence or absence of a species in a specific area (Colvin et al 2015). This is often done by calculating the detection probability of a species in a specified area (or tank) based on quantifiable data such as surveillance surveys, histological surveys, or PCR data (Schmidt et al. 2013; Hunter et al. 2017).

Occupancy models have been implemented to evaluate the performance characteristics of several methods for detecting parasites in wildlife, including serological assays for *Toxoplasma gondii* in arctic-nesting geese (Elmore et al. 2014), to assess test sensitivity and prevalence estimates of *Plasmodium* spp. (Lachish et al. 2011) in wild blue tits (*Cyanistes Caeruleus*), and to estimate detection probabilities of *Schistosoma mansoni* in water (Sengupta et al. 2019). Additionally, our group recently used an occupancy model to evaluate the accuracy of detecting salmon pathogens in histologic sections, including metacercaraie of *Apophallus* spp., *Nanophyetus salmincola*, and the myxozoan (*Parvicapsula minibicornis*) (Colvin et al. 2015). These models provide a powerful tool to quantify detection uncertainty across disparate test systems, allowing for greater accuracy in prevalence estimates.

Here we describe the detection and quantification of *P. neurophilia* DNA in zebrafish tank water using dPCR. The limit of detection was established by spiking water samples with known concentrations of spores, while specificity was verified by testing against *Pleistophora hyphessobryconis*, the only other microsporidium currently recognized to infect zebrafish in a research setting (Sanders et al. 2010). We also evaluated assay performance using various sample types from aquaria holding *P. neurophilia*-infected zebrafish, while implementing a novel multi-state occupancy model to evaluate relationships between habitat, sampling method, distribution, abundance, and detection of parasites in the environment. This newly developed application provides the zebrafish community with a non-lethal diagnostic assay that is both sensitive and specific for *P. neurophilia* using tank water.

## 2. Materials and Methods

### 2.1. Test Development and Optimization

#### 2.1.1. Microsporidian Spore Collection and Purification

*Pseudoloma neurophilia* spores were obtained from hind brain and anterior spinal cords as described by Sanders and Kent 2011. These tissues were mixed with 8 mL of deionized water (diH20) and homogenized using successively smaller-gauged needles (18g, 21g, and 23g). The homogenate was then allowed to sit in a refrigerator at 2° C for 48 hours, while being rigorously vortexed every 24 hours. The homogenate was then centrifuged at max speed (1600g) for 25 minutes. Deionized water was used to hydrolyze host cells, as well as pre-sporogonic *P. neurophilia* stages. The spore suspension was then quantified using a hemocytometer and diluted in (diH2O) as needed to obtain the desired concentrations.

#### 2.1.2 Assay Design & Optimization

Taqman-based qPCR was performed using primers and a probe specific to the ssuRNA gene of *P. neurophilia* include primer and probe sequences here: P10F, P10R, P10Probe) as previously described in Sanders and Kent 2011. Briefly, qPCR was performed in 20 μL reactions composed of 10 μM of forward and reverse primers, 10 μM of hydrolysis probe, 1x TaqMan Universal PCR Master Mix and 2 μL of sample extract using the following reaction conditions: 50 °C for 2 minutes, followed by 40 repetitions of 95 °C for 15 seconds and 60 °C for 1 minute using an Applied Biosystems 7500 Fast Real-Time PCR System and analyzed using the System Sequence Detection Software v1.4.1 (Applied Biosystems).

This assay was then adapted to the droplet digital (ddPCR) platform. ddPCR was performed using DNA extracted from *P. neurophilia* spores collected from zebrafish tank water using a commercial Qiagen DNeasy Blood and Tissue Kit in combination with the forward and reverse *Pseudoloma neurophilia*-specific primers. The reaction was composed of the following rations: 10 μL of supermix for probes – no dUTP (Bio-Rad), 1.8 μL of each primer, 0.5 μL probe (HEX), 1.9 μL water, and 4 μL extracted *P. neurophilia* DNA; total reaction volume = 20 μL. The reaction was run at the Oregon State University Center for Genome and Research and Biocomputing (CGRB) core facilities using a Bio-Rad Qx200 AutoDG Droplet Digital PCR system (Bio-Rad, Hercules, CA) with the following conditions: 10 min at 95 °C followed by 40 cycles of 30 seconds at 94 °C, 1 minute at 60 °C, followed by 98 °C for 10 minutes and held at 4 °C. Individual droplets were then classified as positive (fluorescence present) or negative (no fluorescence) for each reaction using the QX200 Droplet Reader. The number of copies of target DNA present per μL in each reaction was determined using QuantaSoft Analysis Pro software (v1.0.596) by applying Poisson statistics to the ratio of positive droplets to total droplets (Hindson et al. 2011).

#### 2.1.3. Optimization of template DNA input & confirmation of specificity

Sensitivity has been reported to be increased if the initial DNA input is increased in the ddPCR reaction, particularly if the target DNA is in low concentrations (Zhang et al. 2017). Thus, we conducted an additional spiking experiment (as outlined below) at 8,500 spores/liter and extracted DNA from filters using the protocol outlined above. The samples were then prepared for ddPCR with a 2, 3, and 4 μL DNA input in the PCR reaction. The samples were then analyzed using the quantasoft software (Biorad).

Specificity of the original qPCR assay for *P. neurophilia* was established using the Primer-BLAST program (https://www.ncbi.nlm.nih.gov/tools/primer-blast/index.cgi) (Sanders and Kent 2011). This was further validated bioinformatically by performing BLAST searches across several common fish microsporidia, including *Glugea anomala* and *Pleistophora hyphessobryconis*. Specificity in the present study was further validated by testing cross reaction with *P. hyphessobryconis* spores. Spores of *P. hyphessobryconis* spores were collected from zebrafish (*casper* strain) donated from a population known *P. hyphessobryconis* infections and purified in diH20 and treated as described above to obtain an inoculum. Negative control system water was then spiked with 10,000 and 50,000 spores/L concentrations, filtered, and evaluated with DNA extraction and subsequent ddPCR for absolute quantification as described below.

#### 2.1.4. Optimizing the Sonication Protocol

Earlier results indicated that sonication using a probe sonicator for five minutes at a voltage of 55W and a frequency of 20 kHz was the most efficient mode of disrupting the spores to obtain quantifiable DNA from environmental samples (Sanders and Kent 2011). To increase throughput and minimize the potential for cross contamination, a sonicating water bath system, Bioruptor Pico (Diagenode, Denville, NJ), was used. To determine the optimal sonication time required to adequately disrupt spores of *P. neurophilia* a time series was performed using water (negative control tank water) spiked with 8,500 spores/L and processed using the water filtration and DNA extraction protocol described above. The filters were dissolved, submerged in 100 μL PBS, and sonicated at 4, 5, 6, 7.5, 9, 10, and 12.5 minutes using a 30 sec on/off cycle to determine the optimal sonication time.

#### 2.1.5. Sensitivity

Incoming (parasite-free) supply water for the vivarium (dechlorinated city water) was spiked with various spore concentrations for sensitivity testing. The concentration of *P. neurophilia* spores was determined through a series of two separate dilution trials and the limit of detection was first determined using our optimized assay. Two ten-fold serial dilutions were used for experimental exposure: the first from 50 donor fish resulting in a 10^5^, 10^4^, 10^3^, and 10^2^ spores/L series and the second from 40 fish resulting in a 77.5 X 10^4^, 7.75 X 10^3^, 7.75 X 10^2^, 7.75 X 10^1^, and 7.75 spores/L series (Fig. 1; Supplemental Table 2). One L of negative control tank water was filtered and then the filters were spiked with the respective spore concentrations. The filters were then processed for DNA extraction and analyzed via ddPCR. A comparison of sensitivity between qPCR and ddPCR was also conducted utilizing the 10^5^-10^2^ dilution series.

**Figure 1.**
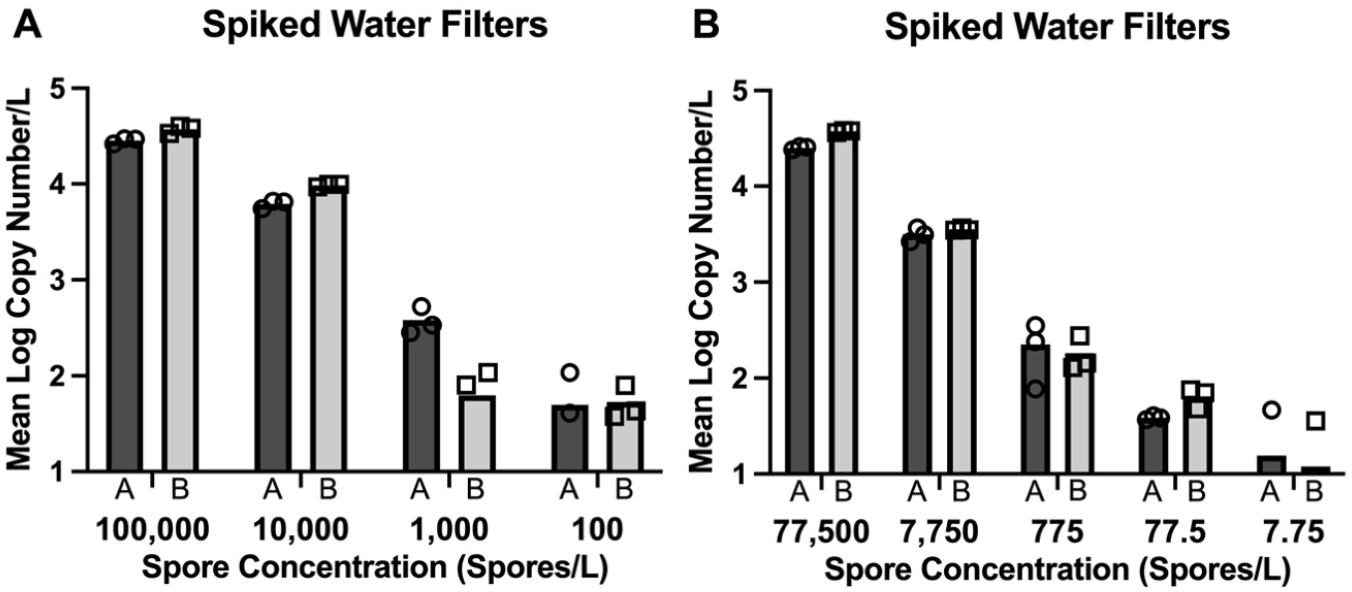
Sensitivity of *P. neurophilia* ddPCR by evaluation of spike filters in duplicate. Two separate experiments: A, with starting concentrations of 100,000 spores/L or 77,500 spores/L. Each DNA assay was conducted in triplicate, with copy number reported for each replicate at each concentration. Circles represent triplicate PCR reactions for filter A, and squares represent the triplicate PCR reactions for filter B.

#### 2.1.6. Water filtration & DNA Extraction

Water samples for all experiments were processed as follows. Replicate 1 L samples of water were filtered as previously described using a vacuum apparatus and a 0.45 μm nitrocellulose filter membrane (Nalgene, Rochester, NY) (Norris et al. 2020). The filter was allowed to dry overnight in an open 15mL conical tube. Once dry, 7mL of acetone was added and vortexed until the filter was completely dissolved. The 15 mL conical tube was then centrifuged for two minutes at 3,000 g. The acetone was removed, and the pellet was resuspended in 1 mL of acetone. 0.5 uL of the solution was then transferred to a 0.65 uL Bioruptor tube and contents pelleted, which was completed twice to accommodate all 1mL of acetone and set in a fume hood to dry. Once all of the acetone had evaporated, 100 uL of 1x PBS was added to the 0.65 mL bioruptor tube and was placed in a Bioruptor Pico, set at 4°C for 18 cycles (9 minutes) with 30 seconds on and 30 seconds off intervals. Once complete, the sample was processed using the Qiagen DNeasy Blood and Tissue Kit (Qiagen, Hilden, Germany) protocol for tissues and eluted in a total volume of 100 μL.

### 2.2. Evaluation of Tanks with Infected Zebrafish

#### 2.2.1. Tank Assembly & Population Dynamics

We further evaluated our assay by testing water from tanks with infected zebrafish. Five tanks of infected fish were tested at three monthly intervals with infected zebrafish held in 16 L tanks containing 30 adult fish/tank. The infected fish originated from five different populations housed at the Zebrafish International Resource Center, Eugene Oregon, and were donated to us following detection of *P. neurophilia* by their staff veterinarian, Dr. Katrina Murray. Populations varied in both sex composition and age. Whereas they were separate populations, a few fish were mixed amongst some tanks so that the experiment would start with the same number of fish/tank. One tank of control (negative 5D line zebrafish fish) was included, originating from the Sinhubber Aquatic Resource Center (SARL), Oregon State University, which was established as a *P. neurophilia* free facility in 2007 (Kent et al. 2011; Barton et al. 2016). The water system is a flow through system with an inflow of 140 ml/min/tank. Water was maintained at 26-28 °C, with conductivity was maintained around 115-130 micro-siemens, and fish were fed once daily with a 1:1 mix of Gemma 300 (Skretting, Westbrook, ME) and Tetramin (Tetra, Melle, Germany). All live animal studies were conducted at Oregon State University and all zebrafish researchers and platform staff were accredited as animal experimentation users according to the FELASA guidelines.

#### 2.2.2. Flow/Static/Spawn Water

Experimental and control tanks were set up and water samples were taken during a flow through period, a static period, and in a static group spawn setting with three monthly sample times. Before collecting water, tank water was mixed by using a scrub brush, stirring tank bottom detritus, and brushing the inside tanks sides. Two liters of tank water were then collected from each tank, filtered through a 0.45 μm filter, and were placed in a −27°C freezer for future DNA extraction. Sampling methods were performed consecutively. Flow sampling was measured first, as our facility is set up as a flow-through system. The tanks were then allowed to normalize for a day, and at the end of the day (1700) flow to each tank was then stopped and left static overnight. Two liters of tank water from the static tanks was then collected the following morning (0900) and filtered. The flow was then resumed for the rest of the day (until 1700).

Group spawn tanks were then set up using all the fish in the tank and at the end of the second day (1700 h). Fish were moved from their regular housing tank into the 16 L spawning tanks with false bottoms and left static overnight to allow for a group spawning event the following morning. Two liters of water were then taken from each spawn tank and filtered through a 0.45 μm vacuum filter as described above. Each filter was saved back frozen at – 27 °C for later ddPCR analysis.

For each of the two water samples collected from a tank at a given time, PCR reactions were carried out in triplicate (Table 1). Copy number results for each reaction are provided in Supplemental Table 3. A tank at a given time was scored as positive when either filter was positive. A filter was scored as positive when at least 2 of the 3 replicates showed detection, regardless of the copy number.

**Table 1.**
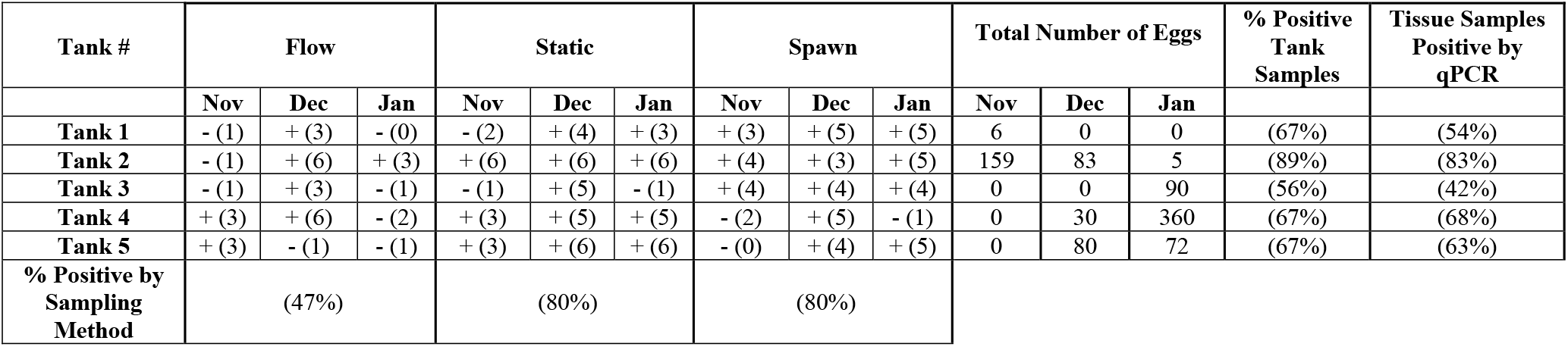
Summary of longitudinal evaluation of five tanks of zebrafish with *Pseudoloma neurophilia* infections. Each cell represents results, positive (+) or negative (-) following ddPCR testing of two 1 L water samples taken from a tank with flowing water (flow), with water turned off for 8 h (static) or water from a group spawn (spawn). DNA extracts from each filter sample were evaluated in triplicate. Number of positive replicates/tanks of the six replicates are in parentheses in each cell (#). A positive score for a tank sample occurred when at least one of the two-liter samples showed detection of two or more replicates. Eggs from successful spawns were collected, counted, and equally divided into 10% pools and tested. Unless the spawns produced a small number of eggs, such as, tank 1 in Nov and tank 2 in Jan, which has a total of 6 and 5 eggs (respectively), thus were grouped into one pool. All egg samples were negative. Neural tissues from fish were evaluated by qPCR after the last water and spawn samples were collected.

| Tank # | Flow |  |  | Static |  |  | Spawn |  |  | Total Number of Eggs |  |  | % Positive Tank Samples | Tissue Samples Positive by qPCR |
| --- | --- | --- | --- | --- | --- | --- | --- | --- | --- | --- | --- | --- | --- | --- |
|  | Nov | Dec | Jan | Nov | Dec | Jan | Nov | Dec | Jan | Nov | Dec | Jan |  |  |
| Tank 1 | - (1) | + (3) | - (0) | - (2) | + (4) | + (3) | + (3) | + (5) | + (5) | 6 | 0 | 0 | (67%) | (54%) |
| Tank 2 | - (1) | + (6) | + (3) | + (6) | + (6) | + (6) | + (4) | + (3) | + (5) | 159 | 83 | 5 | (89%) | (83%) |
| Tank 3 | - (1) | + (3) | - (1) | - (1) | + (5) | - (1) | + (4) | + (4) | + (4) | 0 | 0 | 90 | (56%) | (42%) |
| Tank 4 | + (3) | + (6) | - (2) | + (3) | + (5) | + (5) | - (2) | + (5) | - (1) | 0 | 30 | 360 | (67%) | (68%) |
| Tank 5 | + (3) | - (1) | - (1) | + (3) | + (6) | + (6) | - (0) | + (4) | + (5) | 0 | 80 | 72 | (67%) | (63%) |
| % Positive by Sampling Method | (47%) |  |  | (80%) |  |  | (80%) |  |  |  |  |  |  |  |

#### 2.2.3. Tissue and Egg qPCR

Eggs from each tank were also collected from each spawn, counted, and divided into 10 pools/spawn, with a range of 3 to 58 eggs/pool or if eggs counts < 10, the eggs were pooled in a single sample, as is the case of Tank 1 during the November timepoint (Table 1). At the end of the experiment, each population was euthanized in an ice-bath, and the anterior spinal cord and hind brain tissue from each fish was extracted using fine forceps with the aid of a dissecting microscope. The tissues were then processed as outlined in Sanders et al. 2011. Purified DNA was eluted in 100 μL of buffer AE and extracted DNA from each fish was then quantified using the qPCR assay described above with the same running parameters.

### 2.3. Statistical Analyses

#### 2.3.1 Mixed effects ANOVA

We initially fit a random effects analysis of variance (ANOVA) to partition variation in copy numbers among tanks, replicate water samples within tanks, and replicates within each water sample. We then evaluated differences in copy numbers between the three conditions (flow, static and spawn) and sample month by adding covariates to the ANOVA. The baseline condition in the ANOVA was static and baseline month was December, so parameters should be interpreted relative to the baseline categories. Independence assumptions were evaluated by ordering residuals by tank. The models were fit using R package *lme4* (Bates et al. 2015) implemented in R statistical software version 3.6.1. (R core team 2019).

#### 2.3.2. Multistate Occupancy Modeling

Preliminary results of the ANOVA indicated that the copy counts were too variable among replicates to be reliable indices. However, we believe that they would be useful for classifying counts as none (zero), present, and present and abundant. To evaluate trends and biases in parasite detection classes, we chose to use a multi-state occupancy model (MacKenzie et al. 2009). The multistate occupancy model allows us to evaluate relationships between habitat, sampling method, distribution, abundance, and detection of parasites in the environment. Traditional occupancy models have focused on detection/non-detection data for detection probability estimation; However, in the multistate occupancy model, detection/non-detection is estimated in consideration of three or more states. Thus, we estimated the probability of detecting *P. neurophilia* when it is present in a tank while accounting for treatment effects: flow, static, and spawn water and month. The underlying assumption in this model is that the population is closed, thus occupancy remains consistent during sampling. Here, we considered three states: not detected, detected-but not abundant, detected and abundant. Thereby, our model estimated the following parameters:

Ψ^1^_i, j, k_ = Probability that the parasite is present regardless of abundance;
Ψ^2^_i, j, k_ = Probability that the parasite is abundant given it is present;
p^1^_i, j, k, s_ = Probability that the parasite is detected during sampling occasion, s, given that the true state is present, but not abundant;
p^2^_i, j, k, s_ = Probability that the parasite is detected during sampling occasion, s, given that the true state is present and abundant; and
δ _i, j, k, s_ = Probability that evidence of the abundant state is detected during sampling occasion, s, given that true state is present and abundant,

where i, j, and k denote survey (water sample), tank, and sampling month, respectively. Because of the conditional nature of the probabilities, the probability that a tank contains a large number of parasites (the abundant state is present) is Ψ^1^ * Ψ^2^ and the probability of detecting the abundant state is p^2^ * δ. The abundant state was defined using the raw detection data for each survey from November to December. Based on recommendations from Peterson & Barajas (2018), we calculated the 80^th^ percentile of the maximum copy number across tanks and through time and defined the abundant state as copy numbers that exceeded the 80^th^ percentile.

To implement a multi-state occupancy model, we utilized program Mark (White and Burnham 1999) and *RMark* (Laake 2013) interface package in R statistical software, which uses the conditional binomial version of multistate occupancy to estimate detection probabilities in relation to the three occupancy states. Before fitting the model, we created binary (0, 1) covariates representing: the flow and spawning treatment with static treatment as the baseline category; the samples collected months of November and January with December as the statistical baseline; and tanks two through five with tank one as the baseline. Because the infection status of tanks 1-5 was known, we fixed Ψ^1^ to a value of one (present) and the control tank to a value of zero (absent). We tested for differences between treatments, months, and tanks by fitting a global model containing all of the covariates for the detection parameters (p^1^, p^2^, δ) holding the remaining parameters constant. After model fitting, we retained the statistically significant covariates and the fit the abundant occupancy parameter (Ψ^2^) using all the covariates. A parameter was deemed statistically significant when 95% confidence intervals did not contain zero (i.e., α = 0.05). Only statistically significant covariates for the abundant occupancy parameter were retained in the final model. The occupancy detection parameters can be used to estimate the number of samples required to accurately determine detection of *P. neurophilia* in a tank under each treatment and places the counts from each replicate filter into three categories: not detected, detected – but not abundant, and detected and abundant.

## 3. Results

### 3.1. Test Development

#### Optimization of sonication time

The ideal sonication time was determined based on the reproducibility of replicates and the variability between those replicates (Supplemental Table 1a and 1b). Sonication for 9 minutes (18 cycles) using the Bioruptor Pico was found to be the optimal time to ensure consistently high extraction efficiency of *P. neurophilia* spores in the water with minimal loss of signal.

#### Sensitivity

Two separate experiments were conducted to evaluate the sensitivity of the test using water spiked with log dilutions of known concentrations of spores starting with the highest concentration at either 100,000 or 77,500 spores/L in replicate (Figure 1). Both experiments showed a correlation between DNA copy number and spore concentrations, and results were very similar with filter replicates (A or B) as well as the three replicates for each DNA sample. At lower concentrations, we found that the assay consistently detected down to 100 and 77 spores/L, respectively, with variable results at 7.75 spores/L (Figure 1). Exponential regression values generated from the two spiked filter experiments showed similar trends for detectable copies/spore (Figure 1). For Experiment A, the exponential line of best fit is y = 0.4284 * ln (x) – 0.3442 and for experiment B, y = 0.4188 * ln(x) – 0.2781 (omitting negative reactions). Details of copy numbers at each dilution are reported in Table 2.4. A comparison of sensitivity between qPCR and ddPCR showed that the latter was approximately 1 log more sensitive (Table 1). Furthermore, no inhibition of the reaction was detected when doubling the volume of template DNA from 2 μL to 4 μL at low concentrations of diluted target.

#### Specificity

No DNA detection was observed following testing of filters spiked with a large number of *Pleistophora hyphessobryconis* spores (50,000 or 10,000 spores/L) using the same test parameters as used with *P. neurophilia* samples.

### 3.2. Evaluation of Infected Tanks

#### Consistency between duplicate water samples

A tank was designated positive if *P. neurophilia* DNA was detected in at least two of the three PCR reactions/filter. Results amongst duplicate water samples were consistent (Supplemental Table 3). Out of 45 total tank samples examined, only two tank samples had one 1L replicate testing positive, while the other 1 L sample showed no detection. Likewise, only one of the tank samples showed two replicates from one 1 L being positive while the other 1 L sample had no detection. Data generated from our water test was analyzed using a random effects ANOVA, which found that the majority of variance (88%) occurred between PCR replicates, while the variance between 1-liter samples from the same tank was much less substantial (<1%). The addition of covariates to the ANOVA indicated that the number of copies was significantly lower under the flow and spawning treatments relative to the static treatments (F=47.6, 299 df, p<0.001 and F=23.6, 299 df, p<0.001, respectively). Copies were also significantly lower in November (F=11.5, 302 df, p<0.001) compared to December and January.

#### Flow/Static/Spawn Water

Evaluation of tank water and spawn water by ddPCR at the three-monthly time points showed a range of prevalence amongst the five tanks, with static and spawn water showing higher number of positive detections (Table 1). The five tanks containing infected fish showed a range of prevalence of 56-89%, regardless of sampling method or sample period. Evaluation of data by sample type (flow, static, or spawn) showed that 47% of flow samples were positive, while 80% of the static or spawn water samples yielded were (Table 1). These results were consistent with those from the multi-state occupancy model as discussed below.

#### Multistate occupancy model

The model revealed that detection probabilities, regardless of abundance, was lower in flow samples (0.433) relative to static (0.668) and spawn samples (0.461). When taking abundance into account, detection probabilities for static and spawn increased to (0.85) and (0.90), respectively. Thus, detection probability estimates when the parasite is present but not abundant (Occupancy State 1), were collectively lower than when the parasite is present and abundant (Occupancy State 2). Also, when compared to static and spawn sampling methods, detection was predicted to be lowest when taking samples from tanks with a constant flow (Figure 2). Overall, the detection estimates were much higher when the tank were in an abundant state, however the abundant state was less prevalent during the first month of the survey (November), compared to subsequent months (Table 1). Detection was also lowest in November and January compared to samples collected in December.

**Figure 2.**
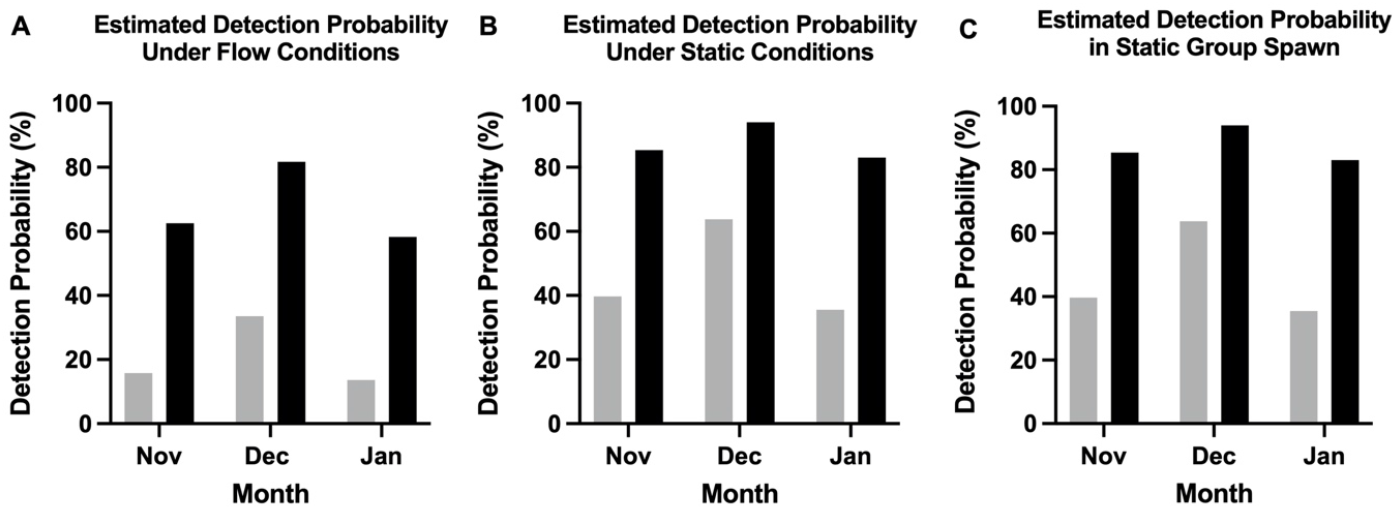
Estimated detection probabilities from the multistate occupancy models by treatment and month for detection of the presence of *Pseudoloma neurophilia*. Probabilities are estimated for detecting the presence of the parasite when the true state is present but not abundant (grey) and present and abundant (black).

#### Fish prevalence compared to tank sample positivity

Prevalence based on PCR of tissues at the end of the 3-month experiment ranged from (42-83%), and the mean value correlated with the number of tanks deemed positive with our water test over the three-monthly samples (r^2^ = 0.85) (Figure 3). For example, in Tank 2, 89% of the tank samples were positive and its fish showed 83% prevalence, whereas Tank 3 had 56% positivity with tank samples and only 42% of its fish were positive (Table 1).

**Figure 3.**
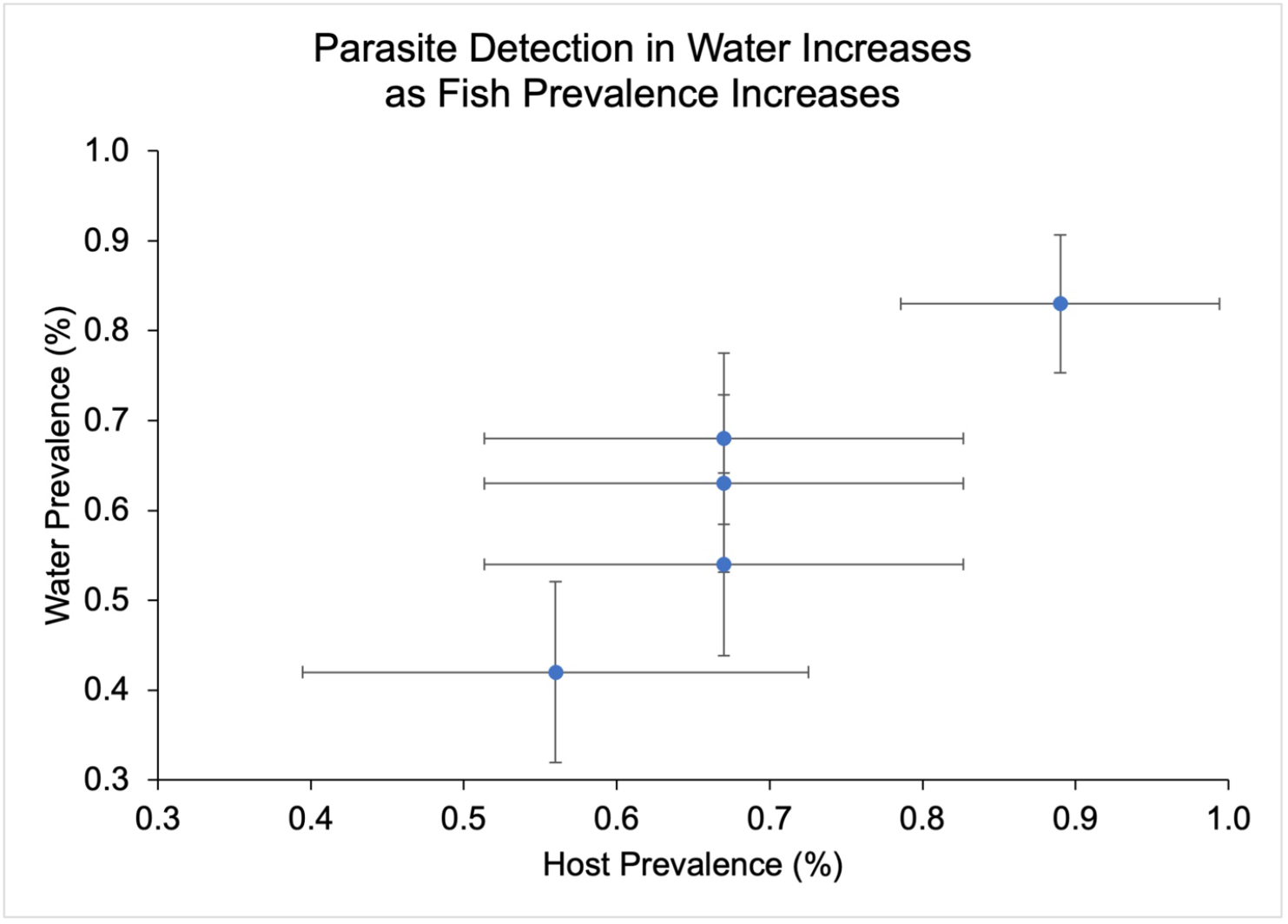
Detection in water increases as prevalence in fish tissues increases. Horizontal error bars = standard error for prevalence amongst the population; Vertical error bars = standard error for water prevalence.

## 4. Discussion

eDNA surveys of fish pathogens are particularly useful for fish kept in captivity, as they are often directed at the population level and defined by a population occupying the same space (e.g., tank). We developed a nonlethal assay that is specific and can provide early detection of *Pseudoloma neurophilia*in zebrafish populations at the tank or facility level. Effective biosecurity programs in modern zebrafish facilities require a combination of approaches, including daily health monitoring and utilization of quarantine rooms to segregate and screen incoming fish. The most common methods to survey for pathogens of zebrafish are lethal and require targeting of specific tissues. Additionally, these assays are usually directed on an individual fish basis, requiring sampling a significant number of subjects to obtain a high confidence estimate of prevalence, abundance, and intensity (Marques and Cabral 2007; Kent et al. 2020b).

Advances in biotechnologies, specifically the development of ddPCR, has distinct advantages for environmental sampling compared to the qPCR method for screening water. Environmental samples often contain humic acid, fulvic acid, as well as debris, which have been shown to negatively impact the performance of DNA-based detection assays (Schrader et al. 2012; Guan et al. 2019). This inhibition can be variable as reflected in a study by Jane et al. 2015, where debris impacted high copy number samples, and in some cases, samples must be diluted to obtain positive results (Schrader et al. 2012; Mckee et al. 2015). The effects of these inhibitory factors are mitigated using ddPCR which divides the PCR into thousands of droplets and analyzes each droplet individually for the target marker. This minimizes the additive effects of inhibitory components in the sample and allows for greater precision when reporting on copy numbers, while also negating the need for a standard curve (Whale et al. 2012; Hayden et al. 2013).

Because the presence of environmental DNA is variable in a given system, sensitivity of eDNA assays is very important, as target DNA in the environment will be dependent on many dynamics of the system (flow rates, stocking density, temperature, etc.) (Jane et al. 2015; Strickler et al. 2015; Troth et al. 2021). Recently, it was demonstrated that fly eDNA was only detectable using a ddPCR based assay when comparing the qPCR and ddPCR assays used to detect Scarce Yellow Sally stonefly (*Isogenus nubecula*) eDNA in water samples (Mauvisseau et al. 2019). Particularly pertinent to the present study, our laboratory also found that a ddPCR test for the nematode *Pseudocapillaria tomentosa* was much more sensitive than qPCR (Norris er al. 2020). Here we observed similar results for the detection of *P. neurophilia* in water (Supplemental Table 4). Combined with pretreatment of samples by sonication, we found that the ddPCR test that we developed for *P. neurophilia* was sensitive and capable of consistently detecting down to 77.5 spores/L.

To our knowledge, this is the first implementation of a multistate occupancy model using the zebrafish host to elucidate trends and biases in parasite detection. Occupancy modeling is typically used in wildlife ecology and has expanded into parasitology for comparison of serological assays in the detection of important pathogens, estimating prevalence, and estimating the probability of detection of important parasites (Lachish et al. 2011; Elmore et al. 2014; Sengupta et al. 2019). Traditional occupancy models have focused solely on detection/nondetection data for detection probability estimations; however, in our multi-state occupancy model, probability estimates were determined using detection/nondetection data with consideration of three states. Thus, we were able to evaluate relationships that may affect the ability to effectively detect *P. neurophilia* in aquaria holding populations of zebrafish (MacKenzie et al. 2009; Peterson et al. 2018).

The occupancy model revealed that detection was lower when tanks were on a constant flow, relative to static and spawn, and that detection was much higher when parasites were predicted to be abundantly present (Figure 2). Detection was estimated to be lowest in November and January, relative to December, likely a consequence due to the timing of exposure. Additionally, microsporidian spores are hydrophobic, which aids in their adherence to host cells (Hayman et al. 2005). This is likely the case with *P. neurophilia*, as we have observed that the spores often accumulate at the air/water interface of bubbles under the coverslip in wet mount preparation. Because of this phenomenon, in combination with the mixing of tank contents during sampling, spores are likely not evenly dispersed in the water column.

Regarding analysis of eDNA samples, replicates for the same DNA extract frequently showed considerable variation. This phenomenon is common with PCR tests at the lower end of detection, and thus triplicate execution of the PCR was run for an accurate estimate of practical repeatability (Ahmed et al. 2009). The variability likely reflecting the variance of detection can be described by the uneven distribution of spores in the tank water, along with forms of parasite DNA (pre-sporgonic stages) being distributed differently between the two separate 1 L samples taken from each tank.

After optimizing the ddPCR assay to screen tank water, we evaluated four sample regimes on non-lethal samples from aquaria with infected zebrafish; water from tanks with flowing water (Flow), water from a tank in which the flow was stopped for 8 h (Static), water from spawning events (Spawn), and eggs from the spawning events. Sensitivity of the assay with static or spawn water samples were similar and detected *P. neurophilia* DNA about 80% of the observed times. In contrast, flowing water samples were positive only 47% of the time (Table 1). The results were similar to earlier studies using a qPCR-based assay (Sanders and Kent 2011), where the parasite was not detected from tanks with flowing water but was detected in spawn water.

*Pseudoloma neurophilia* is commonly found in ovaries, and occasionally in the eggs, thus spores are likely regularly shed into the water during spawning events. In previous studies, *Pseudoloma neurophilia* was detected in water from group spawning tanks, but detection significantly decreased with pair spawns with known infected females (Sanders et al. 2013). In the present study, we consistently detected the parasite in tank water from group spawns, but only at the same rate as static water samples. Whereas *P. neurophilia* is capable of infecting developing eggs, our earlier study showed that the was parasite less frequently detected in embryos than spawn water from the same spawning event (Sanders et al. 2013). Likewise, while water from spawn tanks was frequently positive in the present study, none of the egg pools were positive. Evaluation of prevalence in populations at the end of experiment revealed variable degrees of infection between the fish from different tanks, and the general trend was that tanks with the most infected populations exhibited the highest concentrations of parasite in their respective tanks. The best example of this is in tank 2, which had a prevalence of 83% in fish and had the most consistent water detection compared to the other tanks.

Previous studies have reported that environmental samples are not adequate for the detection of the microsporidium in zebrafish facilities (Crim et al. 2017; Miller et al. 2019). However, this conclusion was based on the sampling of flowing water from aquaria and a sample processing method that did not incorporate a sonication step to disrupt spores as previously recommended (Sanders and Kent 2011). Sonication is a crucial step in detection of spores of *P. neurophilia* as sonicating adequately disrupts the resistant spores making DNA accessible, as seen with *Ovipleistophora ovariae* spores (Phelps 2007). We also demonstrate that sonication improves detection for a better and more consistent result particularly with tank water samples. We were limited to the use of a Diagenode Pico bioruptor, which had a volume limit of 100 μL. Despite this limitation, it is still apparent sonication for 9 min using this sonicating water bath system resulted in the most consistent detection for the microsporidium and thus is a vital element that we recommend for PCR diagnosis of *Pseudoloma neurophilia*, regardless of other methods.

A tank on a given sample day was determined to be positive if at least one of the two 1L water samples resulted in consistent detection of the parasite. We arbitrarily decided that this requires that two of the three DNA replicates were positive from an individual 1L extract.

Although we only evaluated five tanks, there was a correlation with prevalence of infection in fish at the end of the study and number of positive water samples over the three-month period, and this was also supported by a higher rate of detection as determined by the multistate occupancy model. Positivity tended to fluctuate throughout the study, probably reflecting increase of prevalence in fish over time (Ramsay et al. 2009). Future studies coupling current diagnostic methods (histology, whole body PCR, etc.) and this nonlethal water assay would be useful for a longitudinal study to further characterize the infection progression and transmission dynamics. Targeting early onset of infection would elucidate trends regarding the distribution of the parasite in an aquarium setting following initial infection. We are now using this test in a collaboration with the ZIRC, a large zebrafish facility with a history of *P. neurophilia*. With their staff veterinarian (Dr. K. Murray), we are moving forward with screening various populations of fish from their main facility and applying the static water approach to test water from fish following transport. Future studies will focus on coupling this assay with traditional assays (histopathology and tissue qPCR) to investigate the transmission dynamics, particularly targeting the early on-set of infection.

The occurrence of false negatives, in which a water sample was negative from a tank that contained infected fish, demonstrated that the sampling from flowing aquaria reduces sensitivity of the test. The test is still very useful for non-lethal detection of the parasite in its present format with the following recommended applications. Static or spawn water had predicted an occupancy that was always greater than 82% when the parasite was abundant, which was consistently better than predictions under flowing water. Also, static and spawn water detection probability estimates were similar, but the latter would require more effort is it would require setting up spawning tanks/chambers. We therefore recommend leaving tanks static for eight hours with aeration prior to taking a water sample for screening, filtering 1 L water samples following the described extraction protocol.

The positive predictive value of this test is high, as this assay has been proven to be very specific for the microsporidium (Table 2). However, if the test is negative or inconclusive (failed replicates), we recommend tagging the tank as suspect, waiting 24-48 hours, and testing the tank again, as subsequent sampling will increase the probability of detection. For example, with our *P. neurophilia* water test, testing static water twice with independent samples spaced a few days apart increases the sensitivity from 80% to 96% (1 – 0.80 = 0.20 false negative rate). Repeating the test yields 0.20 X 0.20 = 0.04 false negative rate. Hence, 1.0 – 0.04 = a sensitivity of 96 % sensitivity when a second sample is taken a few days later. This approach has commonly been used with other pathogens, in which the test is not optimally sensitive, but the populations to be tested are defined and can be easily retested. In such cases, serial testing has resulted in a near 10% increase in positivity (van Prehn et al. 2015; Larremore et al. 2021).

**Table 2.** Positive (PPV) and negative (NPV) predictive values for each sampling regime by month.

|  | Month | Test Result | Diseased | Not Diseased | Total | PPV or NPV |
| --- | --- | --- | --- | --- | --- | --- |
| FLOW | NOV | Positive | 2 | 0 | 2 | 1 |
|  |  | Negative | 3 | 1 | 4 | 0.25 |
|  | DEC | Positive | 4 | 0 | 4 | 1 |
|  |  | Negative | 1 | 1 | 2 | 0.5 |
|  | JAN | Positive | 1 | 0 | 1 | 1 |
|  |  | Negative | 4 | 1 | 5 | 0.2 |
| STATIC | NOV | Positive | 3 | 0 | 3 | 1 |
|  |  | Negative | 2 | 1 | 3 | 0.33 |
|  | DEC | Positive | 5 | 0 | 5 | 1 |
|  |  | Negative | 0 | 1 | 1 | 1 |
|  | JAN | Positive | 4 | 0 | 4 | 1 |
|  |  | Negative | 1 | 1 | 2 | 0.5 |
| SPAWN | NOV | Positive | 4 | 0 | 4 | 1 |
|  |  | Negative | 1 | 1 | 2 | 0.5 |
|  | DEC | Positive | 5 | 0 | 5 | 1 |
|  |  | Negative | 0 | 1 | 1 | 1 |
|  | JAN | Positive | 4 | 0 | 4 | 1 |
|  |  | Negative | 1 | 1 | 2 | 0.5 |

eDNA survey sensitivity has also been demonstrated to be improved by simply increasing sample volume, sample number, and PCR replicates (Schultz et al. 2015). Another strategy is to target water from older populations of zebrafish, as the infection naturally spreads within a population, resulting in a very high prevalence as fish approach one year of age (Ramsay et al. 2009; Murray et al. 2011). Importantly, when our assay is used with static water, it falls within the new recommendations for assay sensitivity (> 85%), according to the USDA NAHPS (USDA 2021).

## Acknowledgments

This study was supported by the National Institutes of Health under the Research Supplements to promote Diversity in health-related research (NIH ORIP 3 R24 OD010998). We are grateful to the Zebrafish International Resource Center (ZIRC) for providing us with consistent supply of infected subject animals for investigation. We thank Christopher Lawrence, Children’s Hospital Boston, Aquatic Resources Program for providing *Pleistophora hyphessobryconis*. We thank the Sinnhuber Aquatic Resource Center at Oregon State University for providing pathogen free fish. The Oregon Cooperative Fish and Wildlife Research Unit is jointly sponsored by the U.S. Geological Survey, the U.S. Fish and Wildlife Service, the Oregon Department of Fish and Wildlife, Oregon State University, and the Wildlife Management Institute. Any use of trade, firm, or product names is for descriptive purposes only and does not imply endorsement by the U.S. Government. All work was completed at Oregon State University, a land grant university, located on the traditional homelands of the Kalapuya people, who are now absorbed into the Grande Ronde and Siletz Nations.

