## Supplemental Tables for "Multi-state occupancy model estimates probability of detection of an aquatic parasite using environmental DNA: *Pseudoloma neurophilia* in zebrafish aquaria"

**Supplemental Table 1a and 1b**. qPCR values (Ct) from sonication time trials using two serial dilution series to detect *Pseudoloma neurophilia*. (1a) 8,500 spores/L serial dilution series determined that 18 cycles resulted in the best and more consistent detection. (1b) utilizing the 17,000 spores/L serial dilution series analyzed the higher end of sonication times.

| **1a. Sonication Time Trials** | | | |
|---|---|---|---|
| **Concentration** | **Cycles** | **Ct** | **Ct** |
| 8,500 | 8 | 30.94 | 30.83 |
| 8,500 | 8 | 31.73 | 31.33 |
| 8,500 | 10 | 34.13 | 34.64 |
| 8,500 | 10 | 36.03 | 34.49 |
| 8,500 | 12 | 34.16 | 33.89 |
| 8,500 | 12 | 35.05 | 34.95 |
| 8,500 | 15 | 33.84 | 35.09 |
| 8,500 | 15 | 35.34 | 35.07 |
| 8,500 | 18 | 35.22 | 34.06 |
| 8,500 | 18 | 33.62 | 35.58 |
| 8,500 | 20 | 37.6 | 34.12 |
| 8,500 | 20 | 33.69 | 34.07 |
| 8,500 | 25 | 35.02 | 34.52 |
| 8,500 | 25 | 32.87 | 33.24 |
| 850 | 8 | 33.37 | 33.32 |
| 850 | 8 | 34.28 | 35.29 |
| 850 | 10 | 33.11 | 33.95 |
| 850 | 10 | 33.2 | 33.95 |
| 850 | 12 | 33.87 | 33.49 |
| 850 | 12 | 34.78 | 33.84 |
| 850 | 15 | 32.79 | 33.34 |
| 850 | 15 | 32.94 | 33.32 |
| 850 | 18 | 32.03 | 32.95 |
| 850 | 18 | 32.51 | 33.09 |
| 850 | 20 | 32.51 | 32.79 |
| 850 | 20 | 33.21 | 33.08 |
| 850 | 25 | 31.88 | 33.54 |
| 850 | 25 | 35.51 | 34.45 |
| 85 | 8 | 0 | 0 |
| 85 | 8 | 38 | 36.63 |

**Supplemental Table 1a (continued)**

| | | | |
|---|---|---|---|
| 85 | 10 | 37.87 | 0 |
| 85 | 10 | 36.12 | 38.4 |
| 85 | 12 | 0 | 0 |
| 85 | 12 | 36.23 | 0 |
| 85 | 15 | 36.23 | 0 |
| 85 | 15 | 36.99 | 0 |
| 85 | 18 | 36.86 | 37.78 |
| 85 | 18 | 37.62 | 37.09 |
| 85 | 20 | 0 | 35.41 |
| 85 | 20 | 0 | 38.17 |
| 85 | 25 | 0 | 37.08 |
| 85 | 25 | 0 | 37.02 |
| 8.5 | 8 | 0 | 0 |
| 8.5 | 8 | 0 | 0 |
| 8.5 | 10 | 0 | 0 |
| 8.5 | 10 | 0 | 0 |
| 8.5 | 12 | 0 | 0 |
| 8.5 | 12 | 38.2 | 0 |
| 8.5 | 15 | 0 | 0 |
| 8.5 | 15 | 0 | 0 |
| 8.5 | 18 | 0 | 0 |
| 8.5 | 18 | 0 | 0 |
| 8.5 | 20 | 0 | 0 |
| 8.5 | 20 | 0 | 0 |
| 8.5 | 25 | 36.89 | 0 |
| 8.5 | 25 | 0 | 0 |

**Supplemental Table 1b.**

| 1b. Sonication Time Trials | | | |
|---|---|---|---|
| **Concentration** | **Cycles** | **Ct** | **Ct** |
| 17,000 | 10 | 35.15 | 34.52 |
| 17,000 | 10 | 0 | 0 |
| 17,000 | 20 | 33.28 | 33.33 |
| 17,000 | 20 | 32.86 | 32.92 |
| 17,000 | 30 | 32.46 | 32.44 |
| 17,000 | 30 | 32.62 | 32.54 |
| 1,700 | 10 | 37.54 | 36.27 |
| 1,700 | 10 | 36 | 35.85 |
| 1,700 | 20 | 32.69 | 33.01 |
| 1,700 | 20 | 32.56 | 32.38 |
| 1,700 | 30 | 32.88 | 32.64 |
| 1,700 | 30 | 34.98 | 34.56 |
| 170 | 10 | 0 | 0 |
| 170 | 10 | 0 | 0 |
| 170 | 20 | 37.09 | 36.88 |
| 170 | 20 | 37.4 | 36.68 |
| 170 | 30 | 0 | 0 |
| 170 | 30 | 36.99 | 36.54 |
| 17 | 10 | 0 | 0 |
| 17 | 10 | 0 | 0 |
| 17 | 20 | 0 | 0 |
| 17 | 20 | 35.335 | 37.75 |
| 17 | 30 | 0 | 0 |
| 17 | 30 | 0 | 0 |
| 1.7 | 10 | 0 | 0 |
| 1.7 | 10 | 0 | 37.72 |
| 1.7 | 20 | 0 | 0 |
| 1.7 | 20 | 0 | 0 |
| 1.7 | 30 | 0 | 0 |
| 1.7 | 30 | 0 | 0 |

**Supplemental Table 2**. ddPCR results (reported in copy number) used to determine the limit of detection.

| Limit of Detection | | | |
|---|---|---|---|
| **Concentration (Spores/L)** | **Replicates (copies/Rx)** | | |
| 100,000 a | 1188 | 1050 | 1172 |
| 100,000 b | 1596 | 1348 | 1532 |
| 77500 a | 964 | 1042 | 1024 |
| 77500 b | 1528 | 1512 | 1456 |
| 10,000 a | 264 | 260 | 220 |
| 10,000 b | 372 | 394 | 396 |
| 8500 a | 93.6 | 105.4 | 93 |
| 8500 b | 60.6 | 68.4 | 79.2 |
| 7750 a | 125.4 | 146.6 | 105.6 |
| 7750 b | 139.6 | 139.8 | 146.4 |
| 1000 a | 13.46 | 11.28 | 21 |
| 1000 b | 4.32 | 3.22 | 0 |
| 850 a | 11.94 | 5.7 | 3.54 |
| 850 b | 3.64 | 4.8 | 1.778 |
| 775 a | 14.16 | 9.64 | 3.1 |
| 775 b | 11.04 | 5.18 | 5.78 |
| 100a | 4.34 | 1.634 | 0 |
| 100b | 3.18 | 1.516 | 1.73 |
| 85 a | 0 | 0 | 0 |
| 85 b | 0 | 0 | 0 |
| 77.5 a | 1.628 | 1.484 | 1.544 |
| 77.5 b | 2.82 | 1.922 | 3.02 |
| 7.75 a | 1.866 | 0 | 0 |
| 7.75 b | 0 | 0 | 1.432 |

**Supplemental Table 3.** *Pseudoloma neurophilia* tanks with infected zebrafish. Five tanks of infected fish ed at three monthly intervals under three different sampling regimes. Two filters/tank, with triplicate **PCR reactions (reported in copy numbers/µL).**

| **Tank #** | **Filter** | **November** | | | | | | | | |
|---|---|---|---|---|---|---|---|---|---|---|
| | | **FLOW** | | | **STATIC** | | | **SPAWN** | | |
| **1** | **a** | 0 | 0 | 0 | 0 | 0 | 1.8 | 0 | 2.1 | 0 |
| | **b** | 0 | 0 | 1.33 | 0 | 0 | 1.6 | 0 | 3.42 | 1.7 |
| **2** | **a** | 0 | 0 | 0 | 6.78 | 6.02 | 13.1 | 4.94 | 3.24 | 2.22 |
| | **b** | 0 | 1.64 | 0 | 4.8 | 6.6 | 11.3 | 0 | 0 | 1.72 |
| **3** | **a** | 0 | 1.63 | 0 | 1.87 | 0 | 0 | 1.62 | 2.08 | 4.92 |
| | **b** | 0 | 0 | 0 | 0 | 0 | 0 | 0 | 1.86 | 0 |
| **4** | **a** | 1.8 | 0 | 0 | 2.8 | 0 | 0 | 0 | 0 | 3.78 |
| | **b** | 2.92 | 1.81 | 0 | 3.36 | 3.78 | 0 | 3.18 | 0 | 0 |
| **5** | **a** | 3.28 | 0 | 0 | 1.51 | 0 | 2.8 | 0 | 0 | 0 |
| | **b** | 0 | 3.08 | 3.24 | 0 | 2.1 | 0 | 0 | 0 | 0 |
| **NTC** | **a** | NA | NA | NA | NA | NA | NA | NA | NA | NA |
| | **b** | NA | NA | NA | NA | NA | NA | NA | NA | NA |
| **Tank #** | **Filter** | **December** | | | | | | | | |
| | | **FLOW** | | | **STATIC** | | | **SPAWN** | | |
| **1** | **a** | 3.38 | 1.67 | 0 | 4.62 | 1.66 | 3.08 | 0 | 3.18 | 1.69 |
| | **b** | 0 | 0 | 1.62 | 0 | 3.94 | 0 | 3.28 | 1.51 | 3.18 |
| **2** | **a** | 1.82 | 1.66 | 0.3 | 5.1 | 3.68 | 1.66 | 1.67 | 1.86 | 0 |
| | **b** | 1.75 | 1.82 | 1.56 | 11 | 7.22 | 4.54 | 0 | 0 | 1.69 |
| **3** | **a** | 0 | 0 | 0 | 14.2 | 9.72 | 5.9 | 1.69 | 1.73 | 0 |
| | **b** | 1.67 | 1.71 | 4.08 | 3.72 | 6.84 | 0 | 0 | 5.14 | 3.36 |
| **4** | **a** | 7.32 | 2.08 | 1.61 | 4.14 | 4.74 | 4.3 | 0 | 4.72 | 1.88 |
| | **b** | 3.18 | 2.08 | 1.61 | 0 | 1.8 | 4.5 | 5.24 | 4.96 | 4.68 |
| **5** | **a** | 0 | 0 | 0 | 12.2 | 4.82 | 9.16 | 2.92 | 0 | 3.24 |
| | **b** | 0 | 1.8 | 0 | 4.1 | 2.98 | 8.48 | 1.84 | 3.94 | 0 |
| **NTC** | **a** | 0 | 0 | 0 | 0 | 0 | 0 | 0 | 0 | 0 |
| | **b** | 0 | 0 | 0 | 0 | 0 | 0 | 0 | 0 | 0 |

**Supplemental Table 3 (continued)**

| Tank # | Filter | January | | | | | | | | |
|---|---|---|---|---|---|---|---|---|---|---|
| | | FLOW | | | STATIC | | | SPAWN | | |
| 1 | a | 0 | 0 | 0 | 0 | 1.58 | 0 | 0 | 1.59 | 4.6 |
| | b | 0 | 0 | 0 | 0 | 1.63 | 1.74 | 3.34 | 4.22 | 1.63 |
| 2 | a | 0 | 0 | 0 | 15.8 | 19 | 25.4 | 0 | 1.43 | 1.39 |
| | b | 3.62 | 3.44 | 3 | 5.54 | 7.54 | 6.64 | 1.84 | 3.22 | 7 |
| 3 | a | 0 | 0 | 1.63 | 0 | 5.44 | 0 | 1.94 | 0 | 0 |
| | b | 0 | 0 | 0 | 0 | 0 | 0 | 1.52 | 1.44 | 1.45 |
| 4 | a | 1.59 | 0 | 0 | 1.63 | 1.92 | 3.64 | 0 | 0 | 0 |
| | b | 0 | 0 | 3.4 | 0 | 1.72 | 5.18 | 0 | 0 | 1.58 |
| 5 | a | 0 | 0 | 0 | 1.65 | 6 | 1.72 | 1.32 | 0 | 3.16 |
| | b | 0 | 1.71 | 0 | 1.82 | 1.69 | 4.96 | 1.56 | 8.14 | 4.44 |
| NTC | a | 0 | 0 | 0 | 0 | 0 | 0 | 0 | 0 | 0 |
| | b | 0 | 0 | 0 | 0 | 0 | 0 | 0 | 0 | 1.51 |

**Supplemental Table 4.** Sensitivity Comparison: qPCR vs ddPCR. + indicates that the sample had a positive detection by given PCR method with both PCR replicates resulting in detection; - indicates that the sample was either negative or equivocal. Serial dilution series: 100K (0), 10k (-1), 1k (-2), 100 (-3) spores/L.

| | **qPCR** | | | | **ddPCR** | | | |
|---|---|---|---|---|---|---|---|---|
| | 0 | -1 | -2 | -3 | 0 | -1 | -2 | -3 |
| **Detection** | + | + | + | - | + | + | + | + |
